# Placebo-induced pain reduction is associated with inverse network coupling at rest

**DOI:** 10.1101/735563

**Authors:** Isabella C. Wagner, Markus Rütgen, Allan Hummer, Christian Windischberger, Claus Lamm

## Abstract

Placebos can reduce pain by inducing beliefs in the effectiveness of an actually inert treatment. Such top-down effects on pain typically engage lateral and medial prefrontal regions, the insula, somatosensory cortex, as well as the thalamus and brainstem during pain anticipation or perception. Considering the level of large-scale brain networks, these regions spatially align with fronto-parietal/executive control, salience, and sensory-motor networks, but it is unclear if and how placebos alter interactions between them during rest. Here, we investigated how placebo analgesia affected intrinsic network coupling. Ninety-nine human participants were randomly assigned to a placebo or control group and underwent resting-state fMRI after pain processing. Results revealed inverse coupling between sensory-motor and salience-like networks in placebo but not control participants. Specifically, networks were centered on the bilateral somatosensory cortex, as well as on the brainstem, thalamus, striatal regions, dorsal and rostral anterior cingulate cortex, and the insula, respectively. Across participants, more negative between-network coupling was associated with lower individual pain intensity as assessed during a preceding pain task, but was unrelated to expectations of medication effectiveness in the placebo group. Altogether, these findings provide initial evidence that placebo analgesia affects the intrinsic communication between large-scale brain networks, even in the absence of pain. We suggest a model where placebo analgesia increases activation within a descending pain-modulatory network, segregating it from somatosensory regions that might code for painful experiences.

**Highlights:**

- Placebo analgesia affects resting-state connectivity between networks.
- Salience-related and somatosensory regions are negatively coupled at rest.
- This coupling is negative following placebo, but not in control participants.
- More negative between-network coupling is related to lower pain intensity.

## 1. Introduction

Placebos can reduce pain by inducing beliefs in the effectiveness of an actually inert treatment (Price et al., 2008). This top-down effect on pain engages brain regions that signal pain anticipation, treatment expectations, and reward-processes linked to pain relief, such as the lateral prefrontal cortex (Wager et al., 2004, 2007; Watson et al., 2009; Petrovic et al., 2010), rostral anterior cingulate cortex (rACC; Petrovic et al., 2002; Zubieta et al., 2005; Eippert et al., 2009), periaqueductal grey (PAG; Petrovic et al., 2002; Wager et al., 2007), and the ventral striatum (Zubieta et al., 2005; Scott et al., 2007; Geuter et al., 2013). During placebo analgesia, these structures are thought to work in concert with regions associated with actual pain perception, including the thalamus, insula, dorsal anterior cingulate (dACC) and somatosensory cortices (Zubieta et al., 2001; Petrovic et al., 2002; Wiech et al., 2010; Bingel et al., 2011). Previous studies demonstrated enhanced placebo-related coupling between the rACC and PAG (Petrovic et al., 2002; Bingel et al., 2006; Wager et al., 2007; Eippert et al., 2009), as well as interactions between the lateral prefrontal cortex, thalamus, and insula (Lorenz et al., 2003; Wager et al., 2007). Thus, connectivity between regions related to pain processing appears to play a role in placebo-induced pain reduction.

The set of regions involved in placebo analgesia spatially aligns with resting-state networks such as the fronto-parietal/executive control and salience networks (Dosenbach et al., 2007; Seeley et al., 2007), as well as with the sensory-motor network anchored on pre- and postcentral gyri, supplemental motor area, and posterior insula (Beckmann et al., 2005; Damoiseaux et al., 2006; Yeo et al., 2011). The fronto-parietal network (FPN, or “executive control network”, ECN) comprises the lateral prefrontal cortex and inferior parietal regions (Vincent et al., 2008; Yeo et al., 2011) and is relevant for cognitive control (Dosenbach et al., 2006, 2007; Cole et al., 2013). The so-called “salience network” (SN; Seeley et al., 2007) is a sub-system of the FPN centered on the anterior mid-cingulate cortex (or dorsal anterior cingulate cortex, dACC) and insula, and is implicated in attention (Downar et al., 2000), negative affective processing (Seeley et al., 2007) and pain perception (Downar et al., 2003; Legrain et al., 2011; Wager et al., 2013; Kucyi and Davis, 2015). Kong and colleagues (2013), for example, demonstrated resting-state functional connectivity between the FPN/ECN and the rACC which was related to pain modulation in a subsequent task (Kong et al., 2013). Tétreault and colleagues (2016) used resting-state at the beginning of a clinical trial with chronic knee osteoarthritis pain patients to identify placebo responders (Tétreault et al., 2016). They found increased whole-brain coupling of the right midfrontal gyrus that predicted pain relief after placebo treatment. Similarly, Sikora and colleagues (2016) found enhanced connectivity between the rACC and the SN at baseline that was associated with greater symptom reduction after placebo treatment in patients with major depressive disorder (Sikora et al., 2016). Thus, resting-state connectivity involving regions of the FPN/ECN and the SN seems to predict individual placebo responses at baseline, prior to placebo treatment, in different patient populations. Crucially, however, how placebos alter intrinsic network interactions in healthy humans is so far unclear. Therefore, we investigated the effects of placebo analgesia on network interactions at rest within a large-scale, healthy participant sample.

Ninety-nine participants were randomly assigned to a placebo or control group and underwent functional MRI (see also Rütgen et al., 2015). Neural activity during a rest period that completed the scan session was compared between groups. To the best of our knowledge, effects of placebo analgesia on resting-state functional connectivity between networks have not yet been investigated. We thus chose an exploratory and predominantly data-driven analysis approach to do so for the first time. This included the identification of common resting-state networks (Beckmann and Smith, 2004) and the quantification of functional connectivity between them (Smith et al., 2013). Based on previous findings regarding baseline connectivity changes that predicted subsequent (placebo-related) pain modulation (see above; Kong et al., 2013; Sikora et al., 2016; Tétreault et al., 2016), we hypothesized changes in connectivity between the FPN/ECN and SN in placebo compared to control participants, as well as interactions with the sensory-motor network. Additionally, we reasoned that connectivity changes should scale with a behavioral pain intensity score that captured placebo-induced pain relief during a preceding task. Results might shed first light on how placebo analgesia affects whole-brain, intrinsic network fluctuations at rest.

## 2. Materials and methods

### 2.1. Participants

One-hundred-and-twenty participants were randomly assigned to a placebo (*N* = 60, 43 females) or control group (*N* = 60, 38 females). Eighteen participants had to be excluded due to non-responding to the placebo manipulation (10), or technical problems (8). This left 102 participants for the analyses reported by Rütgen and colleagues (2015; placebo group: *N* = 49, 36 females, age range = 20-31 years, mean age = 25; control group: *N* = 53, 34 females, age range = 19-38 years, mean = 26). From this sample, we had to exclude 3 participants (placebo group, 1; control group, 2) due to technical problems with the resting-state data recordings. The current analyses thus included 99 participants (placebo group: *N* = 48, 35 females, age range = 20-31 years, mean = 24; control group: *N* = 51, 34 females, age range = 19-38 years, mean = 25). All were right-handed, healthy, had normal or corrected-to-normal vision, and gave written informed consent prior to participation. The study was reviewed and approved by the ethics committee of the Medical University of Vienna (Vienna, Austria).

### 2.2. Study setup and experimental design

This study was part of a larger project investigating the effects of placebo analgesia on pain and pain empathy (Rütgen et al., 2015). In brief, participants completed a pain calibration procedure upon arrival in the laboratory. After this, the placebo group underwent placebo induction consisting of pill intake and conditioning. Participants then entered the MR scanner and completed a pain, as well as an affective touch task (reported elsewhere), followed by the structural scan. The session was concluded with a resting-state period (**Fig. 1AB**).

**Fig. 1.**
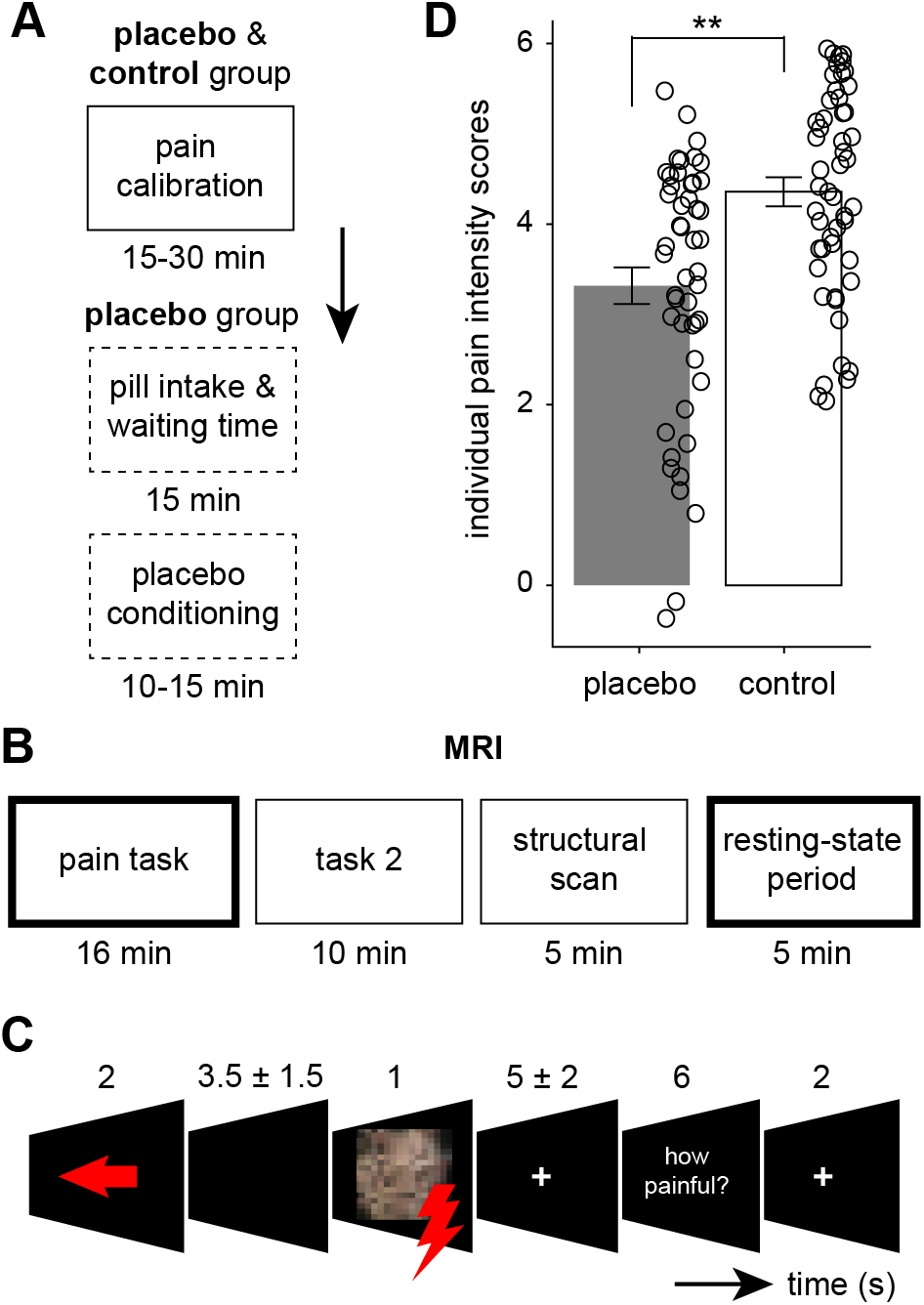
Study timeline, pain task, and individual pain intensity scores. **(A)** Participants of placebo and control groups completed the pain calibration procedure. For the placebo group, this was followed by pill intake, waiting time, and placebo conditioning (dashed framing). **(B)** Tasks and scanning periods that were completed within the MR scanner. Participants first completed the pain task, an affective touch task (task 2, reported elsewhere), the structural scan, and a resting-state period. Bold framing indicates the data used to calculate individual pain intensity scores (pain task; **Section 2.2.4.**), and the data used for the imaging analyses (resting-state period). **(C)** Example for a self-directed, painful trial during the pain task. **(D)** Distribution of individual pain intensity scores in both groups. ** indicates significantly lower pain intensity scores in the placebo compared to the control group, *p* < 0.001.

#### 2.2.1. Electrical stimulation and pain calibration

Individual intensity values (mA) for electrical stimulation were determined during pain calibration (15-30 min). This involved a staircase procedure where participants were asked to rate pain intensity after every electrical shock (500 ms), using a 7-point scale (1, “perceptible, but no painful sensation”; 7, “extreme pain”). The same scale was used for pain intensity ratings throughout the study. Electrical stimulation was delivered using a Digitimer DS5 Isolated Bipolar Constant Current Stimulator (Digitimer Ltd, Clinical & Biomedical Research Instruments) and a bipolar concentric surface electrode attached to the dorsum of the left hand. Shock delivery was controlled manually using Cogent (version 1.32, www.vislab.ucl.ac.uk/cogent.php).

#### 2.2.2. Placebo induction: pill intake and conditioning

Participants of the placebo group were introduced to a medical doctor who administered the placebo pill (starch pill, prepared by the institutional pharmacy of the Medical University of Vienna) and gave information about the alleged medication (see Rütgen et al., 2015 for details). Participants were then asked to rate the expected effectiveness (“Do you expect this medication to be effective in reducing your pain?”; effectiveness_T1_) on a scale from 1 (“not at all”) to 7 (“very effective”). This was followed by a waiting time (15 min) for the alleged medication “to take effect”, after which classical conditioning was used to strengthen the placebo analgesic response (10-15 min; see Rütgen et al., 2015). After this, participants were once more asked how effective they thought the medication was in reducing pain (“How effective is this medication?”; effectiveness_T2_). The control group did not undergo placebo induction and entered the MR scanner after the initial pain calibration (**Fig. 1A**).

#### 2.2.3. Pain task

Inside the MR scanner, participants completed the pain task (16 min), an affective touch task (2 × 5 min, reported elsewhere), and a structural scan (5 min; **Fig 1B**). During the pain task, participants received a cue (2 s) if the electrical shock was directed at themselves (arrow pointing left, self-directed trial) or at another participant (arrow pointing right, other-directed trial). Additionally, the color of the arrow informed the participant about the upcoming stimulation intensity (red, painful; green, non-painful). The other participant was actually a member of the experimental team and never received any shocks. After a brief delay between 2 and 5 s (mean = 3.5 s), participants saw a photo of the shock recipient on the screen (1 s; self-directed trial, scrambled photo of themselves; other-directed trial, photo of the confederate with painful/non-painful facial expression), and a brief electrical shock (500 ms) was delivered (during self-directed trials only). After another fixation period ranging from 3 to 7 s (mean = 5 s), affect ratings (6 s) were collected during one-third of the trials (self-directed pain ratings: “How painful was this stimulus for you?”, other-directed affect ratings: “How painful was this stimulus for the other person?”, and “How unpleasant did it feel when the other person was stimulated?”). Trials were separated with a short fixation period (2 s). In total, participants completed 15 trials per condition (i.e., self-directed painful, self-directed non-painful, other-directed painful, other-directed non-painful). Here, we will focus on self-directed painful and non-painful trials only. An example for a self-directed painful trial is given in **Fig. 1C**. The task was programmed and presented with Cogent (version 1.32, www.vislab.ucl.ac.uk/cogent.php).

Stimulation intensities during self-directed trials for both placebo and control groups were set to calibration-specific stimulation intensities related to pain intensity ratings of 1 (i.e., non-painful trial) or 7 (i.e., extremely painful trial) throughout the pain task. The average stimulation intensities during the pain task were 0.16 ± 0.14 mA (mean ± SEM; pain rating of 1) and 0.74 ± 0.6 mA (pain rating of 6) during nonpainful and painful trials, respectively. Stimulation intensities during non-painful (placebo group: 0.13 ± 0.34 mA, control group: 0.17 ± 0.44 mA: *t*(92.86) = 1.148, *p* = 0.254) and painful trials (placebo group: 0.63 ± 1.36 mA, control group: 0.85 ± 1.91 mA: *t*(89.09) = 1.874, *p* = 0.064) did not differ significantly between groups.

#### 2.2.4. Individual pain intensity scores

We aimed at identifying placebo effects on whole-brain resting-state connectivity. Specifically, we reasoned that placebo effects on between-network connectivity would be related to changes in individual pain intensity ratings. To this end, we leveraged data from the pain task (see above) and calculated a behavioral “pain intensity score” for each participant. We subtracted individual pain intensity during nonpainful trials from pain intensity during painful trials (painful minus non-painful), thereby normalizing individual pain intensity scores for general stimulation intensity. These scores were then correlated with results from the between-network connectivity analysis (**Section 2.4.3.**).

#### 2.2.5. Resting-state period

To assess intrinsic connectivity changes related to placebo analgesic effects, the MR session was concluded with a 5-minute resting-state period. Participants were instructed to remain awake with their eyes open while a white fixation cross was presented at the center of the computer screen. For the placebo group, the resting-state period took place approximately 30 min after the placebo induction (**Section 2.2.2.**). For the control group, the resting-state period took place approximately 30 min after the pain calibration (**Fig. 1AB**).

### 2.3. MRI data acquisition

Imaging data were acquired using a 3 Tesla MRI scanner (Tim Trio, Siemens, Erlangen, Germany) equipped with a 32-channel head coil. We obtained 200 T2*-weighted BOLD images during the resting-state period, using a multiband-accelerated echoplanar imaging (EPI) sequence. Parameters were as follows: TR = 1800 ms, TE = 33 ms, flip angle = 60°, interleaved slice acquisition, 54 axial slices, FOV = 192 × 192 × 108 mm, matrix size = 128 × 128, voxel size = 1.5 × 1.5 × 2 mm. Structural scans were acquired using a magnetization-prepared rapid gradient echo (MP-RAGE) sequence with the following parameters: TR = 2300 ms, TE = 4.21 ms, 160 sagittal slices, FOV = 256 × 256 mm, voxel size = 1 × 1 × 1.1 mm.

### 2.4. Data processing and statistical analysis

#### 2.4.1. MRI data preprocessing

All imaging data were analyzed using the Functional Magnetic Resonance Imaging of the Brain (FMRIB) Software Library (FSL, v5.0.1; https://fsl.fmrib.ox.ac.uk/fsl/fslwiki/; Jenkinson et al., 2012). As a first step, the structural scan was processed (using *fsl_anat)*, including the re-orientation to the Montreal Neurological Institute (MNI) standard space *(fslreorient2std*), bias-field correction (FAST), and brain extraction (BET). The functional images were preprocessed using FEAT. We excluded the first eight volumes to account for T1 equilibration, performed motion correction (MCFLIRT), spatial smoothing with a Gaussian kernel (5 mm full-width at half maximum), and aligned images to the bias-corrected, brain-extracted structural image (FLIRT) using boundary-based registration. The structural image was aligned with the MNI 152 EPI template using non-linear registration (FNIRT). After manual inspection of the registered images, we used independent component analysis (ICA) to automatically detect and remove participant-specific, motion-related artifacts (ICA-based strategy for Automatic Removal Of Motion Artifacts, ICA-AROMA, v0.3-beta; Pruim et al., 2015a, 2015b). Following data de-noising, we applied the registration parameters to the functional data (using *applywarp*).

#### 2.4.2. Group Independent Component Analysis (Group-ICA)

The de-noised functional data were high-pass filtered with a cutoff at 0.01 Hz. We did not apply bandpass filtering as it was shown that high frequencies in the BOLD signal can carry relevant information (Shirer et al., 2015). Data from both placebo and control groups were then submitted to a group-ICA using MELODIC (Beckmann and Smith, 2004). This procedure concatenated all 99 four-dimensional participant data sets into one time × voxel × participant matrix and segregated them into independent spatiotemporal components. The maximum number of independent components (ICs) was set to 30 as this was shown to yield the most common resting-state networks (Abou-Elseoud et al., 2009).

Next, we used a novel, combined approach to separate ICs corresponding to intrinsic resting-state networks from those reflecting noise. We evaluated the 30 ICs in two steps that consisted of manual evaluation (step 1) and an automatic template-matching procedure (step 2). By comparing the ICs to two different, well-known resting-state parcellations (Beckmann et al., 2005; Yeo et al., 2011), we were able to assign the ICs to resting-state networks independent of parcellation-specific network nomenclature.

During step 1, we manually compared each IC to the eight most common resting-state networks as reported by Beckmann and colleagues (2005). This included the (1) medial visual cortical network, (2) lateral visual cortical network, (3) auditory network, (4) sensory-motor network, (5) default-mode network, (6) executive control network, (7) left-, and (8) right dorsal visual networks. Of the 30 ICs, 15 ICs could clearly be assigned to one of the above networks (classified as “signal”), 7 ICs resembled signal-related network fluctuations that could not clearly be assigned to one of the above networks (classified as “unclear/signal”; Griffanti et al., 2017), and the remaining 8 ICs were manually classified as noise-related (“noise”).

During step 2, we used an automatic template-matching procedure (using spatial cross-correlations, fslcc; minimum correlation, *r* = 0.2) to identify ICs that resembled the seven most common resting-state networks as defined by Yeo and colleagues (2011; spatial maps were downloaded from https://surfer.nmr.mgh.harvard.edu/fswiki/CorticalParcellation_Yeo2011). These networks included the (1) ventral and (2) dorsal attention networks, (3) default-mode network, (4) fronto-parietal network (FPN), (5) a network containing motor and auditory regions, (6) a network resembling the salience network (SN), and (7) a network anchored on the ventromedial prefrontal cortex and anterior-ventral temporal lobe. Of the 30 ICs, 22 ICs were assigned to one or more of the above networks (classified as “signal”). The remaining 8 ICs overlapped with ICs that could not clearly be assigned to a specific resting-state network during step 1 (“unclear”). For these, we checked if the ICs were previously classified as “unclear/signal” or “noise”. More specifically, if an IC could not be assigned to a network using spatial cross-correlations (step 2, “unclear”) but was classified as likely being signal-related using the manual evaluation (step 1, “unclear/signal”), then this IC was included in all further analyses. If an IC could not be assigned to a network using spatial cross-correlations (step 2, “unclear”) and if it was classified as noise using the manual evaluation (step 1, “noise”), then this IC was excluded from all further analyses. Overall, 5 out of 30 ICs were classified as noise-related. All remaining analyses were thus performed using a set of 25 ICs (**Fig. 2**). For display purposes, the ICs were brought into 2 mm MNI standard space using FLIRT.

**Fig. 2.**
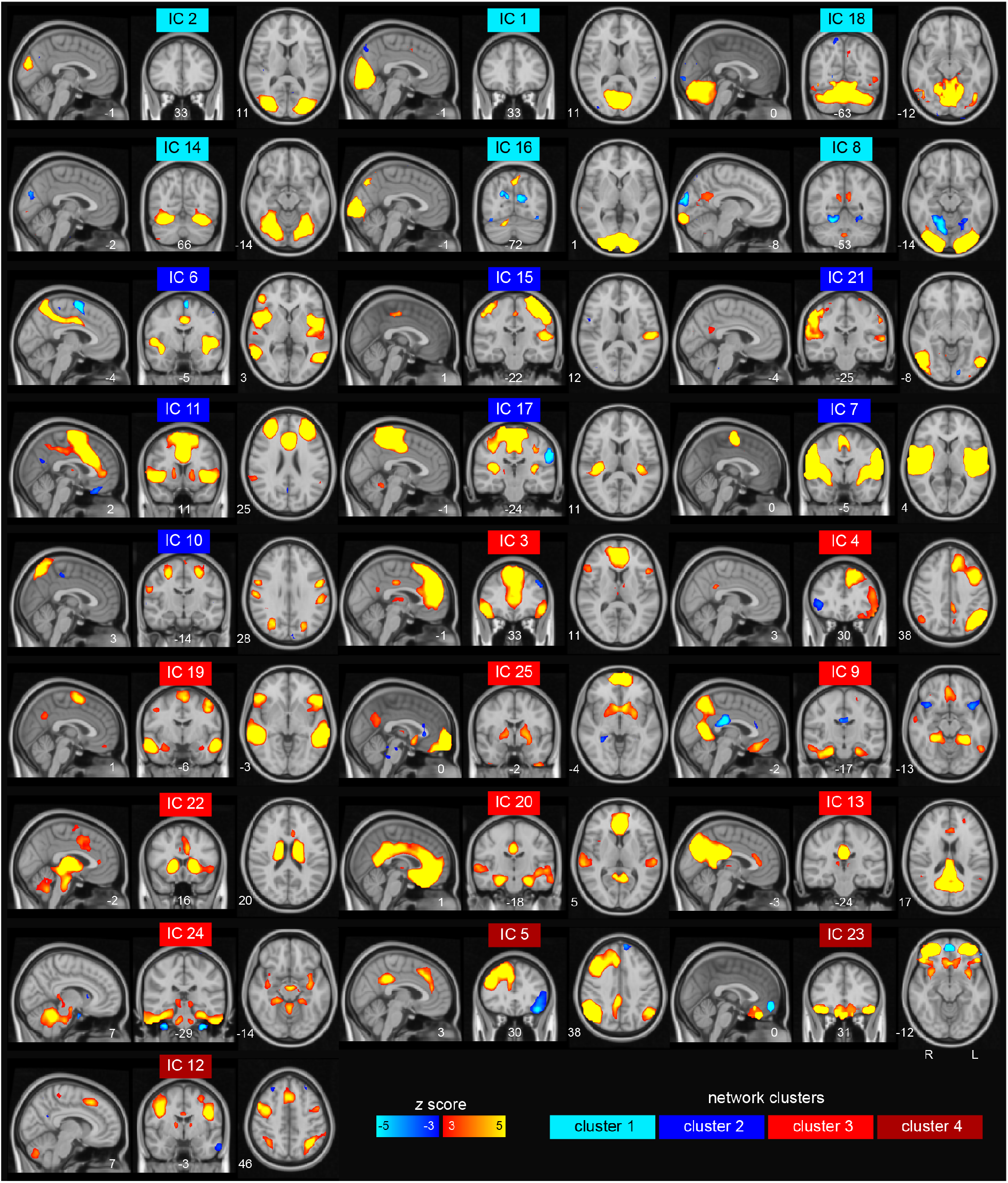
Resting-state networks identified through Group-ICA. Results from group-ICA across both placebo and control groups (*N* = 99) yielded 25 independent components (ICs) that were used for all further analyses. ICs were grouped based on their between-network connectivity which resulted in four clusters of resting-state networks (see also Fig. 3A) centered on visual networks (cluster 1, marked in cyan), sensory-motor networks (cluster 2, marked in blue), networks resembling the DMN (cluster 3, marked in red), and networks resembling the FPN/ECN (cluster 4, marked in dark red). Note that ICs are shown according to neurological convention: R, right; L, left.

After group-ICA, we applied dual regression to generate participant-specific spatial maps and associated time series (Smith et al., 2014) by regressing each IC’s spatial map into the participants’ four-dimensional space × time data set.

#### 2.4.3. Between-network connectivity

Resting-state connectivity between networks was determined using FSL Nets (Smith et al., 2013) and Matlab (Matlab 2017b, The Mathworks, Natick, MA, USA). The participant-specific time series resulting from dual regression (**Section 2.4.2.**) were used to (1) evaluate between-network connectivity across participants, and (2) to compare between-network connectivity between the placebo and control groups. More specifically, we created a 25 × 25 correlation matrix of all ICs (i.e., nodes) per participant where each element represented the correlation strength (i.e., edge) between two nodes. We then calculated full and partial correlation coefficients, the latter to estimate the direct connections between brain regions, and transformed them from Pearson’s *r* to Fisher’s z-values. The participant-specific network matrices were then reshaped into a single row and combined into a participant × edges network matrix to generate the functional connectome of the whole group (i.e., mean network matrix, placebo and control). To group similar nodes, we used hierarchical clustering on the full correlations of the mean network matrix and rearranged the 25 nodes based on their spatiotemporal characteristics using Ward’s method (Ward, 1963; **Fig. 3A**).

**Fig. 3.**
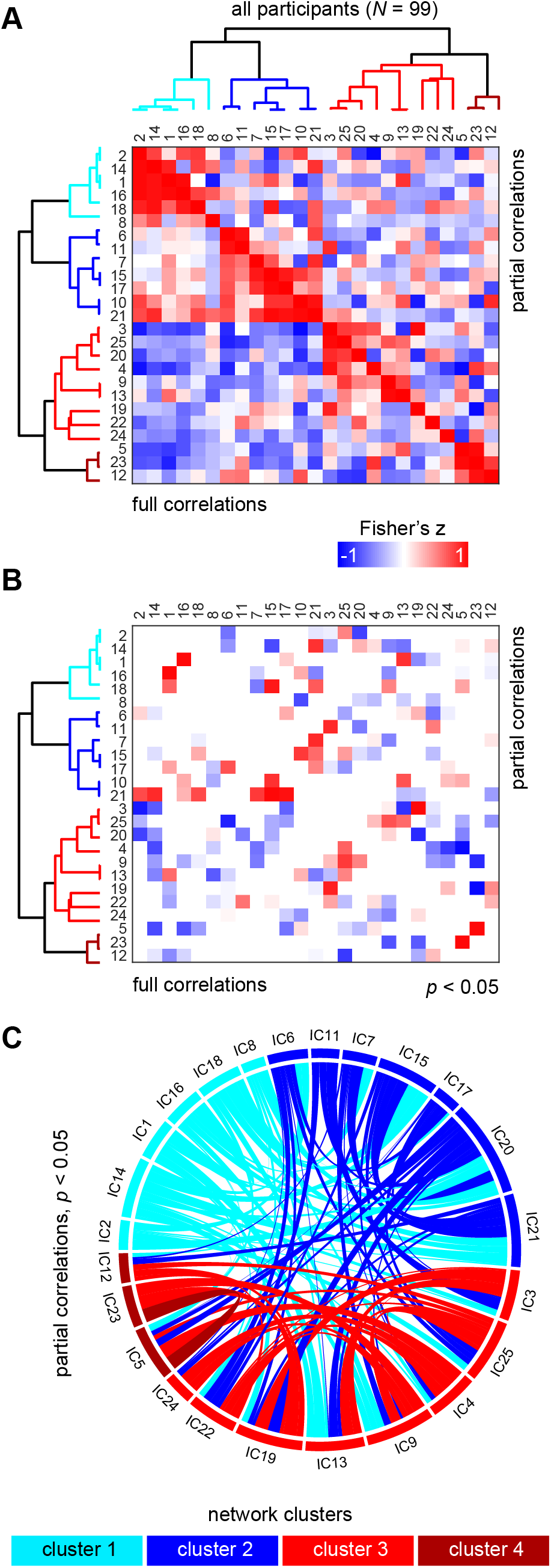
Between-network connectivity across all participants (*N* = 99). **(A)** Hierarchical clustering yielded four groups of resting-state network that were clustered based on their between-network connectivity. The correlation matrix shows full (below diagonal) and partial (above diagonal) correlations between networks. Numeric labels correspond to the independent components (ICs) visible in **Fig. 2**. **(B)** Thresholded connectivity matrix showing only significant between-network connections (*p* < 0.05; corrected for multiple comparisons across all 625 network edges). Significant partial correlations (above diagonal) were used for all further analyses. **(C)** Chord diagram visualizing significant, direct between-network connections across all participants.

Group-level statistics were implemented using FSL’s *randomise* with 5000 permutations per contrast. This tested for a significant effect at each of the 625 network edges by estimating edge-specific p-values, corrected for multiple comparisons across all edges. First, we determined significant between-network connectivity across participants using a one-sample t-test. Second, we assessed significant differences in between-network connectivity between the placebo and control groups using an independent-sample t-test (contrasts: placebo > control, control > placebo). Finally, to test the behavioral significance of the obtained differences in between-network connectivity between groups, we extracted the edge values and performed a cross-participant correlation analysis between individual edge connectivity and pain intensity scores (**Section 2.2.4.**).

#### 2.4.4. Scan-to-scan motion between groups

Lastly, we assured that our connectivity results were not confounded by differences in scan-to-scan motion between the groups. We calculated the temporal derivative time course of the root mean square variance based on all voxels of each brain image (DVARS; Power et al., 2012). This gave an image-specific measure of intensity change by comparing to the previous image. The average DVARS was generally small (mean ± SEM, placebo group: 46.3 ± 1.1; control group: 49.1 ± 1.15) and did not significantly differ between groups (p = 0.081). Thus, the amount of movement during resting-state was comparable between participants of the placebo and control groups.

### 2.5. Data and code availability statement

All anonymized data and analysis code are available upon request in accordance with the requirements of the institute, the funding body, and the institutional ethics board.

## 3. Results

### 3.1. Expectations in medication effectiveness and individual pain intensity scores

To start out, we evaluated the effectiveness of the placebo medication as rated by the placebo group before and after the conditioning procedure (effectiveness_T1_, “Do you expect this medication to be effective in reducing your pain?”; effectivenessT2, “How effective is this medication?”; **Section 2.2.2.**). Ratings significantly increased from effectiveness_T1_ (mean ± standard error of the mean, SEM: 4.35 ± 0.16) to effectivenessT2 (4.75 ± 0.21, t(47) = −2.479, *p* = 0.017, *d* = −0.31, [95% CI, −0.72, 0.1]), indicating that the conditioning procedure increased expectations of medication effectiveness.

To test the effects of placebo on subjective pain intensity, we compared individual pain intensity scores obtained during the pain task between the groups (i.e., pain ratings during painful minus non-painful trials; **Section 2.2.4.**). Pain intensity scores were significantly lower in the placebo (mean ± SEM: 3.32 ± 0.41) compared to the control group (4.36 ± 0.32, t(90.031) = 4.028, *p* < 0.0005, *d* = −0.82, [95% CI, −1.23, −0.4]; **Fig. 1D**). Thus, placebo successfully decreased subjective pain intensity with an effect size similar to previous placebo analgesia neuroimaging studies (Zunhammer et al., 2018).

### 3.2. Between-network connectivity across groups

As a first step, we identified 25 resting-state networks based on data from both placebo and control groups (**Fig. 2, Section 2.4.2.**). Using these, we calculated the mean connectivity between every network (i.e., node) which yielded a correlation matrix of between-network connectivity values (i.e., edges; **Fig. 3A**). Highly correlated nodes were grouped together during hierarchical clustering (**Section 2.4.3.**). This resulted in four network clusters (**Fig. 2 and 3**): *cluster 1* contained visual networks (centered on visual cortex but also on the cerebellum; marked in cyan); *cluster 2* contained sensory-motor networks (anchored on the motor, somatosensory and auditory regions, but also dACC and bilateral insula; marked in blue); *cluster 3* comprised components of the default-mode network (DMN; including frontal and posterior medial regions, the hippocampus, surrounding medial temporal lobe, and parietal regions; marked in red); and *cluster 4* included a right lateralized fronto-parietal/executive control network (FPN/ECN; marked in dark red).

Notably, cluster 3, which was centered on the DMN, also contained a sub-cluster that was only loosely connected to the remaining networks. This sub-cluster included a network that resembled the salience network (SN; IC 22; **Fig. 2**, marked in red). Another network that was spatially similar to the SN (IC 11) showed between-network connectivity with components of cluster 2 which contained sensory-motor networks (**Fig. 2**, marked in blue). While cluster 4 included a right lateralized FPN/ECN (IC 5; **Fig. 2**, marked in dark red), its left lateralized counterpart (IC 4) was mainly coupled with networks of cluster 3. Thus, both FPN/ECN and SN revealed between-network connectivity with networks of different clusters rather than mainly coupling with networks belonging to the FPN.

### 3.3. Group differences in between-network connectivity

Most importantly, we hypothesized differences in between-network connectivity between the placebo and control groups. We expected that these connectivity changes should involve the FPN/ECN and SN, as well as interactions with the sensory-motor network in placebo compared to control participants.

Result revealed connectivity between a network assigned to cluster 2 (centered on sensory-motor networks; IC 15) and a network assigned to cluster 3 (centered on the DMN; IC 22) that differed significantly between the placebo and control groups (control > placebo; *p* = 0.047, corrected for multiple comparisons across all 625 network edges; **Fig. 4A**). These networks were negatively correlated in placebo participants (one-sample t-test against zero: *t*(47) = −5.281, *p* < 0.0005) and not significantly correlated in participants of the control group (*p* = 0.761; **Fig. 4B**). In other words, IC 15 and 22 were inversely coupled in the placebo compared to the control group such that decreased activation within IC 15 was associated with increased activation within IC 22 and vice versa. IC 15 comprised a network including the bilateral somatosensory cortex, supplemental motor area, posterior insula/left superior temporal gyrus, and the cerebellum (**Fig. 4C**, upper panel; **Table 1**). IC 22 consisted of a network between the cerebellum and brainstem, bilateral thalamus and striatal regions, dorsal and rostral ACC, and the anterior insula (**Fig. 4C**, lower panel; **Table 1**). There were no other significant group-differences in between-network connectivity.

**Fig. 4.**
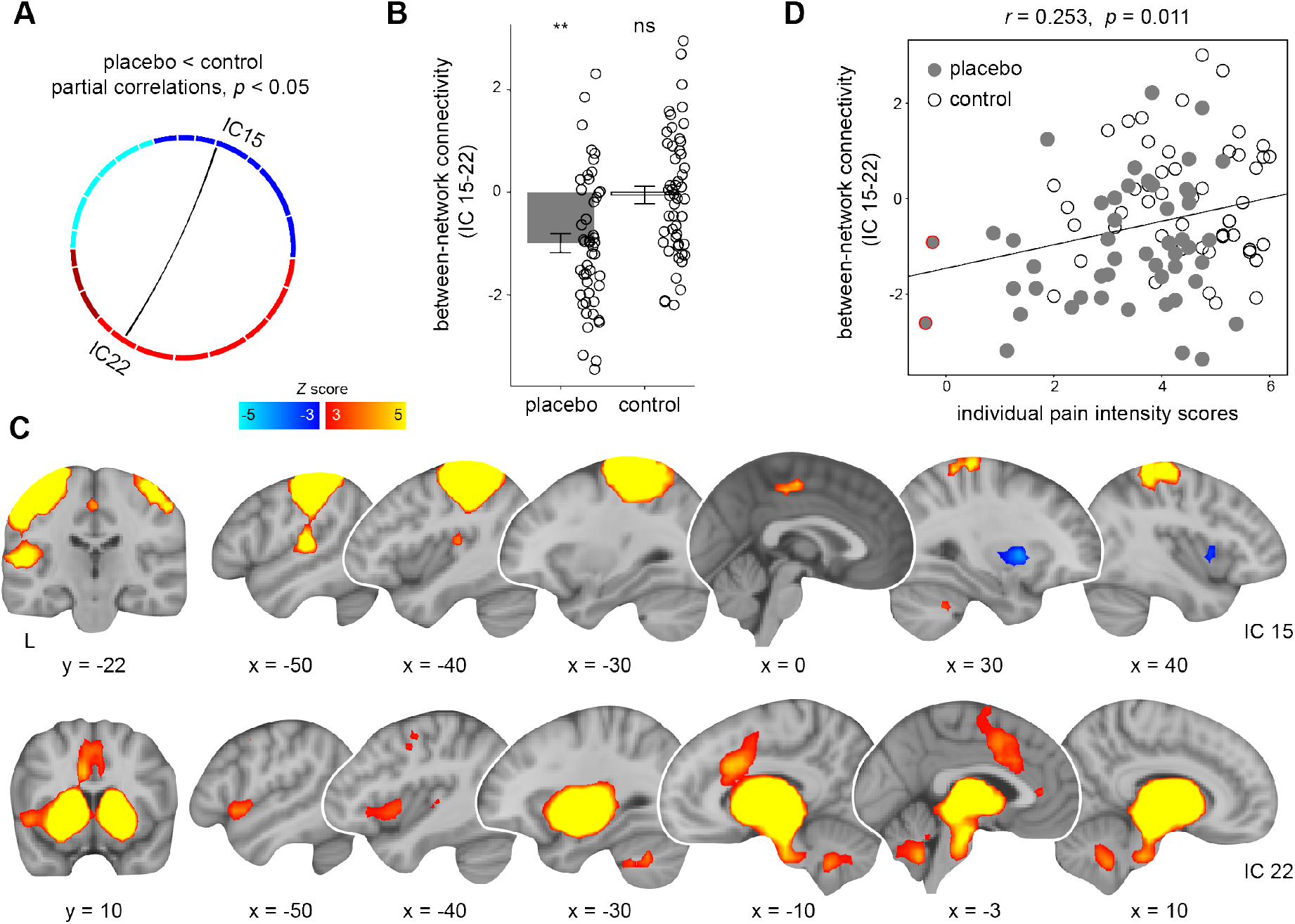
Between-network connectivity differences between placebo and control groups. **(A)** Chord diagram visualizing the connectivity between independent component (IC) 15 and 22 was significantly different in placebo compared to control participants (placebo < control, *p* = 0.047, corrected for multiple comparisons across all 625 network edges). **(B)** Between-network connectivity was negative in the placebo group (**, *p* < 0.001) but not significant (ns) in the control group. **(C)** IC 15 (upper panel, initially grouped together with cluster 2; **Fig. 3A**) and IC 22 (lower panel, initially grouped together with cluster 3; **Fig. 3A**). Coordinates of brain regions are described in **Table 1**. Brain slices are based on the mean structural scan of all participants. L, left. **(D)** Results showed a positive correlation of between-network connectivity (IC 15-22) with individual pain intensity scores across participants (*N* = 99). Thus, more negative coupling was associated with lower subjective pain intensity during the preceding pain task (**Fig. 1CD**). The correlation remained robust when removing two possible outliers from the placebo group (mean - 3 standard deviations, marked in red).

**Table 1.** Networks IC 15 and 22. MNI coordinates represent the location of peak voxels. We report the first local maximum within each cluster. Percentages describe the probability of the peak voxel belonging to the labelled region. Anatomical nomenclature was obtained from the “Harvard-Oxford Cortical Structural Atlas”, the “Harvard-Oxford Subcortical Structural Atlas”, and the “Cerebellar Atlas in MNI152 space after normalization with FNIRT”. IC, independent component; L, left; R, right.

| ICs and brain regions | MNI coordinates |  |  | z-value | cluster size |
| --- | --- | --- | --- | --- | --- |
|  | x | y | z |  |  |
| IC 15 |  |  |  |  |  |
| 51% Postcentral Gyrus, 5% Precentral Gyrus, L | -46 | -22 | 56 | 17.1 | 941 |
| 35% Postcentral Gyrus, 12% Superior Parietal Lobule, R | 42 | -38 | 64 | 7.04 | 210 |
| Cerebellum, R | 18 | -50 | -32 | 6.03 | 90 |
| 60% Precentral Gyrus, 14% Juxtapositional Lobule Cortex (formerly Supplementary Motor Cortex), L | -2 | -18 | 52 | 5.13 | 37 |
| Cerebellum, 58% R IX, 28% Vermis VIIIb, R | 6 | -62 | -44 | 3.26 | 7 |
| 6% Temporal Occipital Fusiform Cortex, 5% Occipital Fusiform Gyrus, L | -26 | -66 | -20 | 3.69 | 5 |
| 1% Lateral Occipital Cortex, inferior division, L | -30 | -82 | -28 | 3.26 | 4 |
| 2% Occipital Fusiform Gyrus, L | -42 | -74 | -28 | 3.15 | 1 |
| IC 22 |  |  |  |  |  |
| 91% Thalamus, L | -14 | -10 | 8 | 18.5 | 2489 |
| Cerebellum, 69% R Crus I, 25% R VI, R | 34 | -54 | -36 | 5.34 | 299 |
| 34% Precentral Gyrus, 2% Middle Frontal Gyrus, L | -42 | -6 | 44 | 3.65 | 20 |

### 3.4. Between-network connectivity and relation to individual pain intensity scores

So far, we reported lower pain intensity scores and inverse coupling between two resting-state networks (IC 15-22) in placebo compared to control participants. As a final step, we tested the cross-participant relationship between behavioral pain intensity scores and between-network connectivity using correlation analysis. Results demonstrated a significantly positive correlation of between-network connectivity (IC 15-22) and individual pain intensity scores across all participants (*r* = 0.253, *p* = 0.011; bootstrapped 95% CI based on 5000 samples = [0.08, 0.44]), that remained stable after the exclusion of two possible outliers (placebo group; *r* = 0.218, *p* = 0.032, bootstrapped 95% CI [0.03, 0.41]; **Fig. 4D**). Altogether, this indicates that greater inverse coupling between networks that were anchored on the bilateral somatosensory cortex (IC 15), as well as on the brainstem, thalamus, striatal regions, dorsal and rostral ACC, and the anterior insula (IC 22) was related to lower subjective pain intensity during the preceding pain task.

### 3.5. Between-network connectivity and relation to expectations in medication effectiveness

To mitigate the effect of treatment expectations on resting-state fluctuations, we assessed connectivity after pain processing took place. However, treatment expectations might have still affected our results. We clarified this by testing the relationship of between-network connectivity and individual expectations in medication effectiveness assessed before (effectiveness_T1_) and after (effectivenessT2) placebo conditioning in participants of the placebo group (Section 3.1.). Results showed no significant correlation of between-network connectivity with expectations in medication effectiveness rated before (effectiveness_T1_: *p* = 0.912) or after placebo conditioning (effectiveness_T2_: *p* = 0.364). Thus, between-network connectivity was not related to expectations in medication effectiveness in the placebo group.

## 4. Discussion

Effects of placebo analgesia on resting-state functional connectivity between networks have not yet been investigated so far. Here, we report first findings from an exploratory and predominantly data-driven analysis approach. We assessed the effects of placebo analgesia on network interactions during rest within a large-scale sample of healthy participants. Results showed significant inverse coupling between two networks that were centered on the bilateral somatosensory cortex (IC 15) and on the brainstem, thalamus, striatal regions, dorsal and rostral ACC, and the insula (IC 22) in placebo compared to control participants. The amount of inverse coupling was positively related to individual pain intensity scores across participants. In other words, more negative between-network connectivity was associated with lower subjective pain intensity during the preceding task. Results were unrelated to expectations of medication effectiveness in the placebo group. Our findings highlight how placebo analgesia affects intrinsic network fluctuations during rest.

We hypothesized network interactions between the FPN/ECN, SN, and the sensory-motor network that should differ between placebo and control groups. Results partly confirmed our expectations as we found negative connectivity of a network that resembled the SN with a somatosensory network in placebo compared to control participants (**Fig. 4**). The classic SN consists of the dACC and insula, and is connected to subcortical regions such as the thalamus, (ventral) striatum, and the PAG (Seeley et al., 2007). The dACC was implicated in attention (Downar et al., 2000) and conflict monitoring (Botvinick et al., 2004), negative affect including anxiety and stress (Seeley et al., 2007; Hermans et al., 2011), pain perception (Downar et al., 2003; Legrain et al., 2011; Wager et al., 2013; Kucyi and Davis, 2015), but also reward-based decision making (Bush et al., 2002; Hayden and Platt, 2010) and positive affective processing (Bartels and Zeki, 2004). In a similar vein, the insula was associated with salience (Menon and Uddin, 2010), affective (Singer et al., 2009) and pain processing (Rodriguez-Raecke et al., 2010; Wiech et al., 2010), as well as interoception (Craig, 2002, 2009). Altogether, the SN was proposed to direct attention to relevant sensory input, to integrate sensory information with internal bodily states, and to prepare appropriate actions (Menon and Uddin, 2010; Gogolla, 2017).

Besides the common SN regions, the network we identified (IC 22) further comprised the rACC which is regarded as central for the down-regulation of subjective pain intensity during placebo analgesia (Wager and Atlas, 2015). Together with the PAG, the rACC is part of a descending circuit for pain modulation that further includes the rostral ventromedial medulla and its projections to the spinal cord (Fields, 2004). Previous studies using human neuroimaging showed placebo-related coupling between the rACC and PAG during pain processing (Petrovic et al., 2002; Bingel et al., 2006; Wager et al., 2007; Eippert et al., 2009), as well as increased connectivity of the rACC with the SN that predicted individual placebo responses at baseline (Sikora et al., 2016). Thus, interactions between the rACC and regions of the SN, including the PAG, appear crucial for placebo-induced pain relief.

The somatosensory network (IC 15) comprised the bilateral somatosensory cortex, supplemental motor area, and left posterior insula/superior temporal gyrus, and we propose that it potentially coded for the somatosensory representation of the noxious stimulation during the preceding task. Connectivity between IC 15 and 22 was negative in placebo compared to control participants such that increased activation in one network was related to decreased activation in the other and vice versa. We suggest that increased activation of brainstem, thalamus, striatal regions, dorsal and rostral ACC, and the insula (IC 22) underlies descending pain modulation after placebo which in turn might be associated with lower activation of somatosensory regions (IC 15). Across participants, the degree of inverse coupling between the networks was associated with lower individual pain intensity scores obtained during the preceding pain task (**Fig. 4D**), probing the behavioral relevance of these resting-state network fluctuations. This is in line with findings from Petrovic and colleagues (2002) who demonstrated similar engagement of rACC and brainstem in both placebo and opioid analgesia. Increased activation of brainstem, thalamus, striatal regions, dorsal and rostral ACC, and the insula (IC 22) might thus reflect placebo-induced, opioidergic activation related to the top-down modulation of pain (Baumgärtner et al., 2006; Eippert et al., 2009; Zunhammer et al., 2018). Most importantly, we found these connectivity changes during resting-state, in absence of actual pain anticipation or perception. Our findings thus suggest that placebo analgesia might affect communication between large-scale networks, potentially up-regulating opioidergic signaling even when pain is not present.

Network connectivity implies oscillatory synchrony between neuronal assemblies which is thought to facilitate information processing (Buzsaki and Draguhn, 2004). Certain brain networks, such as the FPN and the DMN, are often negatively correlated during resting-state (Fox et al., 2005; but see for example van Buuren et al., 2019) as well as during cognitively demanding tasks (Cole et al., 2012; but see Douw et al., 2016), whereby the magnitude of inverse coupling was linked to better task performance (Anticevic et al., 2012; Cole et al., 2012). Thus, negative between-network connectivity appears to affect information transfer in a functionally relevant manner. Fox and colleagues (2005) suggested that while positive connectivity might be involved in the integration of information across different brain regions, negative connectivity might segregate neuronal processes that subserve opposing goals. Negative coupling between the SN and somatosensory network might thus indicate increased activation of descending pain-modulatory processes after placebo, segregating this network from somatosensory activation that might code for (preceding) noxious stimulation.

We did not find interactions of SN and somatosensory network with the FPN/ECN. A reason for this might be that we assessed resting-state functional connectivity after pain processing took place. Previous studies that investigated placebo effects on network connectivity focused on baseline resting-state periods in order to predict pain modulation (Kong et al., 2013) or individual placebo responses (Sikora et al., 2016; Tétreault et al., 2016). For instance, Kong and colleagues (2013) found increased resting-state coupling between the FPN/ECN and the rACC to be related to pain modulation in a subsequent task. The lateral prefrontal cortex is considered as central node of the FPN (Vincent et al., 2008; Yeo et al., 2011), important for cognitive control (Dosenbach et al., 2006, 2007; Cole et al., 2013) and the top-down modulation of pain (Wager et al., 2004, 2007; Watson et al., 2009; Petrovic et al., 2010). Previous results reporting FPN/ECN engagement were thus potentially related to treatment expectations or pain anticipation. We propose that our results are unrelated to such factors, as we measured resting-state fluctuations after pain processing took place. Furthermore, there was no significant relationship of between-network coupling and expectations in medication effectiveness across participants of the placebo group.

One possible explanation of our results could be that activation changes from the preceding task persisted into resting-state, thereby inducing the observed connectivity effects. If this was the case, we would assume comparable task-based activation changes, leading to significant connectivity between SN and somatosensory networks in both groups. Contrary to this, we did not find significant connectivity between the SN and somatosensory network in the control group (**Fig. 4B**). Thus, while participants showed inverse network coupling following placebo, we found network *decoupling* in the control group. Following this line of thought, several limitations need to be addressed. First, although we found significant differences in between-network connectivity between placebo and control groups, interpretation is limited by the between-subject nature of the study design. To further investigate placebo-induced changes in intrinsic network fluctuations, future studies should measure resting-state during placebo as well as control conditions in the same participants. Resting-state periods before and after placebo induction and pain processing might further be helpful to disentangle the effects of treatment expectations, pain anticipation and pain processing on network coupling, and to investigate trait characteristics that render individuals more or less susceptible to placebo-related belief inductions (Vachon-Presseau et al., 2018). Second, we found that inverse network coupling was positively related to individual pain intensity scores which we obtained from the preceding pain task (**Fig. 1C**). Future studies should use a more in-depth assessment of pain to generate individual pain intensity scores based on multiple tasks in order to replicate the current findings. However, in spite of these limitations, we provide first evidence for placebo-induced network changes during resting-state and their relation to behavioral pain modulation after pain processing.

## 5. Conclusions

To conclude, we investigated placebo effects on whole-brain resting-state functional connectivity in healthy human participants after pain processing. Our results revealed inverse coupling between two networks in placebo but not control participants. These networks were anchored on the bilateral somatosensory cortex, as well as on the brainstem, thalamus, striatal regions, dorsal and rostral ACC, and the anterior insula, respectively. Across participants, more negative between-network coupling was associated with lower individual pain intensity as assessed during a preceding pain task, but was unrelated to expectations of medication effectiveness in the placebo group. Altogether, these findings provide initial evidence that placebo analgesia affects the intrinsic communication between large-scale brain networks, even in the absence of pain. Our findings suggest that placebo analgesia increases activation within a descending pain-modulatory network, segregating it from somatosensory regions that might code for painful experiences.

## Acknowledgements

The authors would like to thank Daniel Graf, Bernadette Hippmann, Julia Hebestreit, Alexander Kudrna, and Andreas Martin (all University of Vienna) for assistance with functional MRI measurements; Fritz Zimprich (Medical University of Vienna) for medical support; and Andreas Gartus (University of Vienna) for technical support. The study was supported by the Vienna Science and Technology Fund (WWTF; Project CS11-016).

